# CD4 T Cells Acquire Cytotoxic Properties to Modulate Cellular Senescence and Aging

**DOI:** 10.1101/2024.01.14.575313

**Authors:** Yehezqel Elyahu, Ilana Feygin, Noa Pinkas, Alon Zemer, Amit Shicht, Omer Berner, Ekaterina Eremenko, Anna Nemirovsky, Keren Reshef, Lior Roitman, Valery Krizhanovsky, Alon Monsonego

## Abstract

Aging is characterized by the progressive deterioration of tissue structure and function, leading to increased vulnerability to diseases and, eventually, death. One prominent process in aging is the accumulation of senescent cells. Although the immune system has been recognized as crucial for the elimination of senescent cells, the associated mechanisms remain incompletely understood. Here we show that CD4 T cells differentiate into cytotoxic T lymphocytes (CTLs) in a senescent cell-rich environment and that a reduction in the senescent cell load, achieved using chemical senolytic drugs, was sufficient to halt this differentiation. We further demonstrate that eliminating CD4 CTLs in the context of late aging by selectively deleting the Eomes transcription factor in CD4 T cells resulted in the increased accumulation of senescent cells, profound physical deterioration, and a decreased life span. In liver cirrhosis, a model of localized chronic inflammation, CD4 CTL elimination increased the senescent load and worsened the disease. Collectively, our findings demonstrate the fundamental role of CD4 CTLs in modulating tissue senescence and unveil a new aspect of age-related T-cell biology implicated in disease susceptibility and longevity.

## Introduction

A fundamental characteristic of aging is the gradual deterioration of nearly all tissues ^1,2^, leading to decreased functionality and increased vulnerability to age-related diseases. At the cellular level, the decline in tissue integrity during aging has been associated with the accumulation of senescent cells (SCs) ^3^ along with stem cell exhaustion ^1^ and reduced immune competence ^4,5^. Cellular senescence is a terminal state that has been implicated as a core cause of various age-related diseases, including cardiovascular diseases ^6,7^, dementia ^7,8^, and cancer ^9,10^. This cellular state can be induced by DNA damage, oncogenic mutations, mitochondrial dysfunction, mechanical stress, or infections ^11^. SCs gradually populate aged tissues ^12,13^ and are characterized by cell cycle arrest ^11^, resistance to apoptosis ^14^, and the secretion of pro-inflammatory cytokines, referred to as senescence-associated secretory phenotype (SASP), often causing long-lasting inflammation which disrupts tissue hemostasis ^15,16^.

The modulation of tissue senescence is largely mediated through interactions with the immune system ^17^. Both innate and adaptive immune mechanisms have been implicated in the process of SC elimination ^17–19^. While CD8 T cells and NK cells are associated with SC clearance ^20,21^, the intricate interplay between SCs and immune cells remains ambiguous, especially in the context of aging, which is characterized by the marked impairment of many immune functions ^22,23^. We recently found that CD4 CTLs accumulate in aging, comprising up to 50% of the CD4 T cell compartment in old mice ^24^. These cells are characterized by the production of cytotoxic molecules such as granzyme B (GzmB) and perforin that are classically produced by CD8 T cells, T helper 1 (Th1)-related cytokines such as TNF and IFN-γ, and chemokines including CCL3, CCL4, and CCL5 ^17,25^. The main transcription factor (TF) implicated in their differentiation is eomesodermin (Eomes) ^26^, which has been shown to be essential for the production of these lytic molecules and cytokines ^17,25^. In line with our findings, a human study focused on a small cohort of supercentenarians observed similar unexpected frequencies of CD4 CTLs, which were associated with the well-being and longevity of these individuals ^27^. Others have shown that in aged humans, CD4 CTL accumulation increases in the context of chronic diseases such as multiple sclerosis ^28,29^. Moreover, a recent study by Hasegawa et al. found that human cytomegalovirus (CMV) reactivation in skin senescent fibroblasts induced CD4 CTL recruitment and cytotoxicity against SCs ^30^. However, a causal relationship between the senescent environment, the differentiation of CD4 CTLs, and their impact on SC load at the tissue and organism levels has not been elucidated to date.

Here, we show that CD4 CTLs differentiate in an environment rich in SCs and play a key role in modulating tissue SC abundance. Targeted depletion of Eomes-positive CD4 T cells in localized senescence models and aged mice resulted in worse disease outcomes, accelerated aging, and reduced lifespan. These findings have the potential to contribute to novel immunotherapies for treating age-related diseases and enhancing physical well-being during aging.

## Results

### CD4 T cells differentiate into cytotoxic cells in a senescent-rich environment

The cellular landscape of the immune system profoundly changes with aging ^24^. Specifically, within the CD4 T cell compartment, we and others have reported the marked accumulation of effector, exhausted, Tregs, and cytotoxic cells in both aged mice and humans ^24,27^. However, it remains largely unknown whether these changes result from the extrinsic age-dependent molecular milieu or stem from cell-intrinsic alterations. To address this question, we adoptively transferred 20 million leukocytes from young (2- to 3-month-old) CD45.1 mice to either young (2-3 months; young→young) or old (22-24 months; young→old) CD45.2 mice. After 1 month, we collected spleens from these young and old CD45.2 mice and analyzed the frequencies of CD4 T cell subsets positive for CD45.1 (young-transferred cells) and CD45.2 (young or old endogenous cells) using flow cytometry (Fig. 1a, Extended Data Fig. 1a, Methods). Strikingly, compared to the young→young group, in the young→old group we observed a significant increase in the percentage of CD45.1^+^ CTLs (CD3^+^CD8^-^CD4^+^EOMES^+^CCL5^+^) and exhausted CD4 T cells (CD3^+^CD4^+^CD44^+^PD1^+^, Fig. 1b-c, Methods). In contrast, the frequencies of other CD4 T cell subsets such as naïve cells (CD3^+^CD4^+^CD62L^+^CD44^-^), effector cells (CD3^+^CD4^+^CD62L^-^ CD44^+^PD1^-^), and Tregs (CD4^+^CD3^+^FOXP3^+^) did not change significantly (Fig. 1d, Extended Data Fig. 1b, Methods) between the young and old environments. Next, we compared the proportions of CTLs and exhausted cells out of total CD4 T cells in the young-transferred versus old-endogenous cells in mice from the young→old group. Notably, the young-transferred cells reached similar proportions of both CD4 CTLs and exhausted cells as those endogenously present in these old mice (Fig. 1e), suggesting that their proportions highly depend on environmental cues.

**Figure 1.**
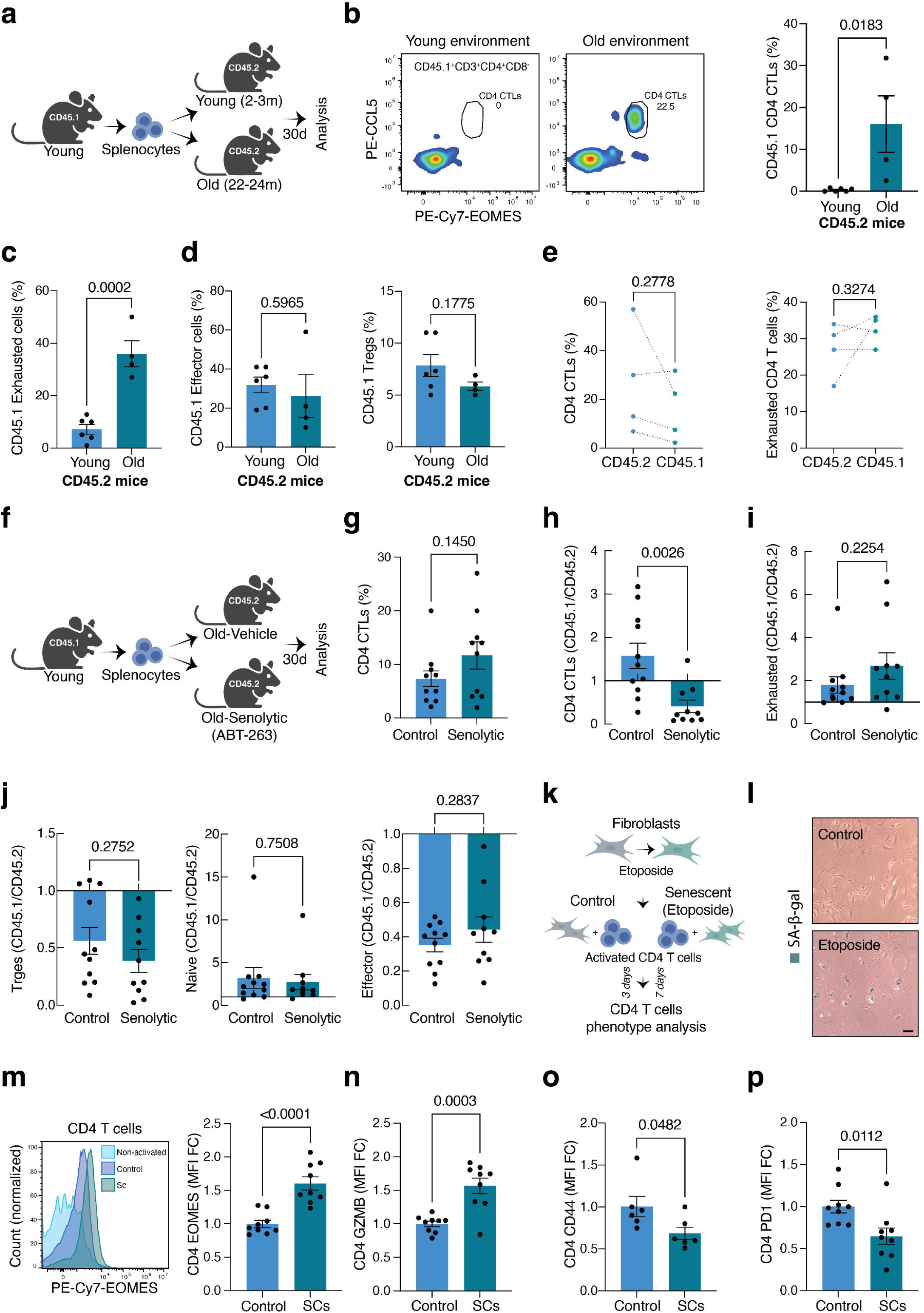
Adoptively transferred young-origin CD4 T cells differentiate into CD4 CTLs in an aged environment *in vivo* and in the presence of senescent cells *in vitro*. **a.** Experimental setup: splenocytes were harvested from young CD45.1 mice, and then 20 million of these cells were injected into either young (2-3 months) or old CD45.2 (22-24 months) mice, followed 30 days later by the harvesting of spleens for analysis. **b**. Left: Representative flow cytometry plots illustrating the proportions of CD4 CTLs (EOMES^+^CCL5^+^) out of CD4 T cells (CD3^+^CD4^+^CD8^-^) among transferred cells (CD45.1^+^) in young or old CD45.2 mice. Right: Quantitative analysis of CD4 CTLs among transferred cells in young (n=6) or old (n=4) WT mice (CD45.2). **c.** Quantitative analysis of exhausted cells (CD3^+^CD4^+^CD62L^-^CD44^+^PD1^+^) among transferred cells in young (n=6) or old (n=4) WT mice (CD45.2) **d.** Quantitative analysis of effector cells (CD3^+^CD4^+^CD62L^-^ CD44^+^PD1^-^) and T regulatory cells (Treg; CD3^+^CD4^+^FOXP3^+^) among transferred cells in young or old WT mice (CD45.2). **e.** The percentages of CD4 CTLs (Left) and exhausted cells (Right) out of CD4^+^ cells. Comparison between CD45.1 (young-transferred cells) and CD45.2 (old-endogenous cells) in the old group. **f.** Experimental setup: splenocytes were harvested from young CD45.1 mice, and then 20 million of these cells were injected into old mice treated with vehicle (10% ethanol, 30% PEG 400, 60% Phosal50) or Navitoclax (ABT-263, 50 mg/kg,) for 30 days, after which spleens were harvested for analysis. **g.** Quantitative analysis of CD4 CTLs among CD45.2 (old-endogenous cells) cells in control (n=11) and senolytic drug-treated mice (n=10). **h-i.** The ratio of CD4 CTLs **(h)** or exhausted cells **(i)** originating from CD45.1 (young-transferred) and CD45.2 (old-endogenous) cells in control (n=11) and senolytic drug-treated mice (n=10). **j.** The ratios of effector cells (Left), naïve cells (CD3^+^CD4^+^CD62L^+^CD44^-^, Middle), and Treg (Right) cells originating from CD45.1 (young-transferred) and CD45.2 (old-endogenous) cells in control (n=11) and senolytic drug-treated mice (n=10). Each dot represents the ratio in a single mouse. **k.** Experimental setup: fibroblasts derived from mouse lungs were treated with etoposide to induce senescence. Both control untreated (Control) and senescent fibroblasts (SCs) were then co-cultured with polyclonally activated young CD4 T cells (2-3 months old) for 3 or 7 days and then CD4 T cells were collected for flow cytometry analysis. **l.** Representative images of senescence-associated β-Galactosidase (SA-β-Gal) staining in control (upper panel) and etoposide-induced senescent fibroblasts (lower panel). Scale bar represents 50 μm. **m.** Left: a representative flow cytometry histogram plot illustrating the expression of EOMES in CD4 T cells harvested from non-activated control (light blue), activated with control fibroblasts (dark blue), or activated with senescent fibroblasts (green), after 7 days of co-culture. Right: quantitative analysis of EOMES median fluorescence intensity (MFI) of activated CD4 T cells co-cultured with either control or senescent fibroblasts. **n-p.** MFI analysis of granzyme B (GZMB) **(n)**, CD44 **(o)**, and PD1 **(p)**. Each dot represents an individual technical well. For each experiment, CD4 T cells were purified by negative selection from 7 mice (**m-p**). Bars represent mean ± SEM from one (**o**), two (**a-e, k-p,** or three (**f-j**) independent experiments. Data were analyzed using a two-tailed Student’s t-test (**a-p**). Exact P-values are presented in the figures.

To specifically evaluate the contribution of SC load to the differentiation of CTLs and exhausted cells, we transferred young CD45.1 cells into old CD45.2 control mice (Vehicle) or old CD45.2 mice treated with the senolytic drug ABT-263 (Navitoclax) for one month prior to cell transfer (Fig. 1f). Compared with control animals, the mice treated with senolytic drug displayed a reduction in their SC burden as represented in the liver tissue (Extended Data Fig. 1c-d). One month following the transfer, we analyzed the young-transferred and old-endogenous cells in these mice using flow cytometry. First, we assessed the percentage of endogenous CD4 CTLs in the spleens of Vehicle- or senolytic drug-treated mice. We observed no difference in the frequency of endogenous CD4 CTLs in the senolytic drug-treated group (Fig. 1g). Remarkably, when we compared the ratio of young-transferred to old-endogenous CD4 CTLs in both the control and senolytic drug-treated groups, this ratio was lower in the senolytic drug-treated group (Fig. 1h), indicating reduced differentiation towards CD4 CTLs after senolytic treatment. Notably, the senolytic treatment did not impact the ratio (young-transferred versus endogenous cells) of other CD4 T cell subsets, including exhausted cells, Tregs, naïve cells, and effector cells (Fig. 1i-j). Together, these results demonstrate a causal link between aging-related SC accumulation and the differentiation of CD4 T cells into CD4 CTLs. To further establish this connection, we polyclonally stimulated young CD4 T cells in the presence of either etoposide-induced senescent fibroblasts or control fibroblasts for 3 and 7 days (Fig. 1k, Methods). The induction of senescence was validated via senescence-associated β-galactosidase (SA-β-gal) staining, which was positive in the etoposide-treated but not in the control fibroblasts (Fig. 1l). At the two tested time points, we noted a significant increase in the expression of EOMES in the CD4 population co-cultured with etoposide-treated fibroblasts compare to control (Fig. 1m, Extended Data Fig. 1e). Moreover, CD4 T cells co-cultured with the senescent fibroblasts significantly increased their expression of GzmB compared to the cells from the control group (Fig. 1n, Extended Data Fig. 1e), along with reduced expression of CD44 and PD1 (Fig. 1o-p, Extended Data Fig. 1f), suggesting an inhibitory effect of SCs on CD4 T cell activation.

### CD4 CTL depletion in aged mice is associated with increased senescent cell load, frailty, and decreased survival

To examine the impact of CD4 CTLs on aging phenotypes and lifespan, we generated a mouse model that allows for the conditional depletion of CD4 CTLs. We and others have previously identified the Eomes TF as a key element involved in the differentiation of CD4 CTLs ^24,27,31,32^. To downregulate this TF specifically in CD4 T cells, we bred CD4-Cre^ERT2^ mice with Eomes-floxed mice to create CD4-Cre^ERT2+/-^Eomes^fl/fl^ mice which were then administered with tamoxifen (TMX). To validate our model’s efficacy and specificity, we treated young CD4-Cre^ERT2-/-^Eomes^fl/fl^ (Control) or CD4-Cre^ERT2+/-^Eomes^fl/fl^ mice (Eomes-KO) with subcutaneous injections of TMX (Methods). We then activated spleen-derived T cells using anti-CD3/anti-CD28 coated beads for 24 hours and analyzed EOMES expression by T cells via flow cytometry. Whereas no differences were observed in the CD8 T cell subset, we observed a significant reduction in EOMES-expressing CD4 T cells and EOMES expression within CD4 T cells in Eomes-KO as compared to control mice (Extended Data Fig.2a-b, Methods). Additionally, we did not observe changes in CD4 molecule expression between groups or differences in IL-2 or IFN-γ production, measured in cell supernatants, suggesting that EOMES depletion in CD4 T cells does not affect their activation and/or cytokine production (Extended Data Fig. 2c-d, Methods).

Utilizing our model, we evaluated physical performance and SC load in 20-month-old mice before and after TMX exposure for 45 days (Fig. 2a, Methods). While frequencies of CD4 CTLs (out of all CD4 T cells) were increased in the blood of control mice, they were reduced in Eomes-KO mice (Fig. 2b-c). These results strengthen our previous observation that CD4 CTLs accumulate with aging ^24^ and validate the *in vivo* depletion of Eomes-expressing CD4 T cells. To assess physical performance, we first employed the endurance test to evaluate both groups before (day 0) and after (day 45) TMX treatment. By comparing the performance ratio (post-treatment/pre-treatment) for each mouse, we observed a marked decline in the Eomes-KO group as compared to the control group (Fig. 2d). For a more in-depth physical evaluation, we monitored multiple physical parameters in both groups using metabolic cages (Extended Data Fig. 3a-b), before and after 45 days of exposure to TMX-containing food, and then compared the performance ratios (Methods). No physical differences were observed between the groups before treatment (Extended Data Fig. 3c-g). During the experiment, we recorded a higher incidence of death in the Eomes-KO than in the control group (Extended Data Fig. 3h). Strikingly, we detected a reduction in activity (measured as the sum of fine and directed movements in meters, Fig. 2e-f, Methods) and increased food consumption (Fig. 2g) in the Eomes-KO group relative to the control group. The water intake remained consistent between the two groups (Fig. 2g).

**Figure 2.**
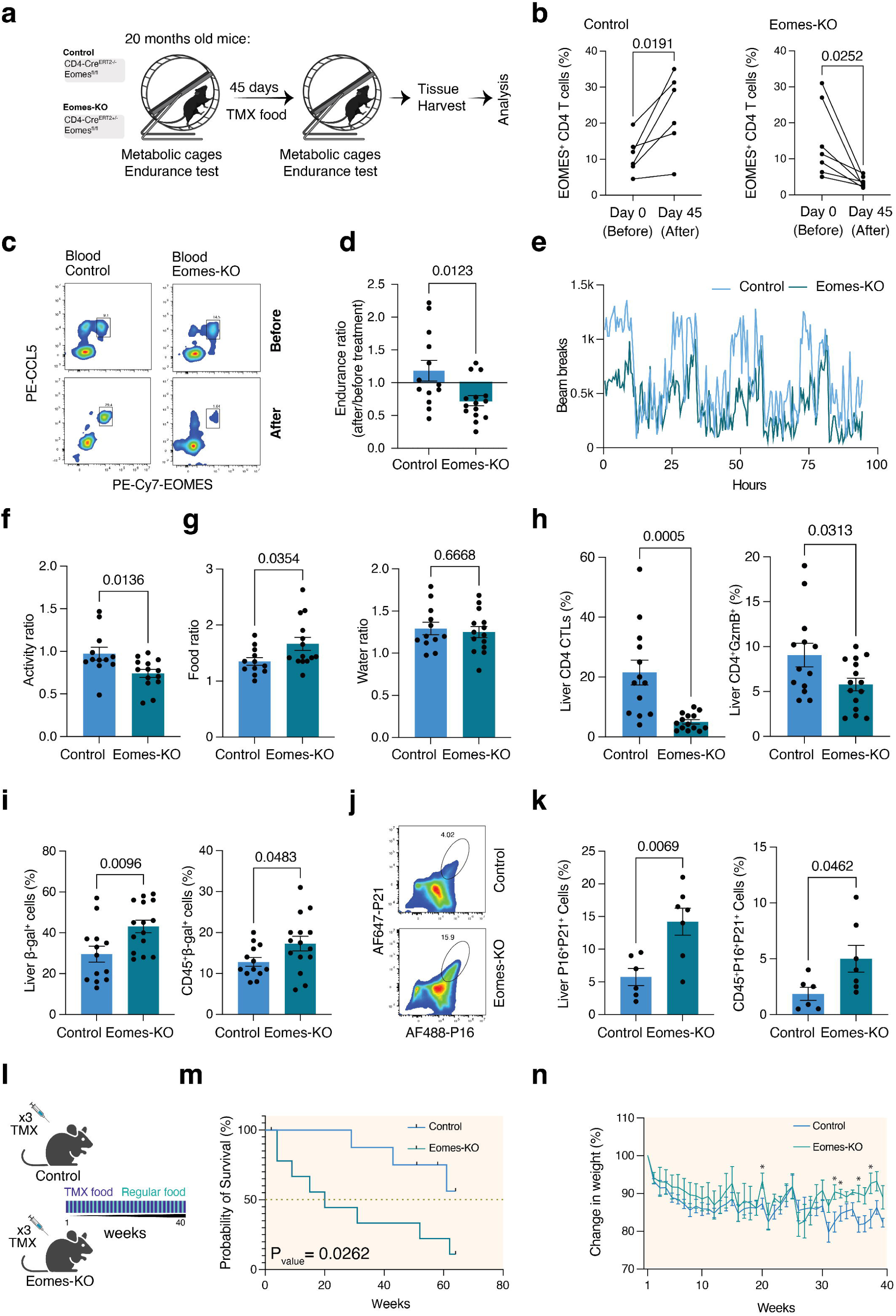
Decreased physical performance and survival in aging are associated with lower frequencies of CD4 CTLs. **a.** Experimental setup: the physical performance of aged (20 months old) control (Cre^ERT2-/-^Eomes^fl/fl^; Control) and CD4 CTL-depleted (Cre^ERT2+/-^Eomes^fl/fl^; Eomes-KO) mice was evaluated using the hanging test and metabolic cages. Additionally, 100-200 µl of blood was collected from the tail vein of each mouse for flow cytometry analysis. Subsequently, the mice were administered a TMX regimen, which included intraperitoneal injections of 100 µl of TMX for 3 days, followed by an alternating dietary regimen of two weeks on TMX chow and two weeks on regular chow, lasting for a total of 6 weeks. After the TMX exposure period, physical performance was reevaluated in each mouse. Finally, the mice were sacrificed, and blood, spleen, and liver samples were collected for further analysis. **b.** Graphs showing CD4 CTLs (CD3^+^CD4^+^EOMES^+^CCL5^+^) in the blood before and after TMX treatment in the control (Left, n=6) and in the Eomes-KO (Right, n=7) groups. **c.** Representative flow cytometry plots showing CD4 CTLs in the blood of control (Left) or Eomes-KO mice (Right), before (Upper plot) and after (lower plot) TMX treatment. The X-axis represents EOMES expression, and the Y-axis represents CCL5 expression. **d.** Hanging test results as a ratio between scores after and before TMX for each mouse in the control (n=13) and Eomes-KO (n=15) groups. Each dot represents a single mouse. Scores were normalized to the body weight of each mouse (Methods). **e.** Representative beam breaks (Y-axis) recording for control (n=4) or Eomes-KO (n=4) mice from 4-day recordings with a 30-minute interval resolution. **f-g.** The ratio between post-treatment and pre-treatment performance in metabolic cages for each mouse in both the control (n=12) and Eomes-KO (n=14) groups. Each dot represents the ratio for one mouse. **f.** Activity ratio, which represents the sum of fine motor movement and intentional movements in the control and Eomes-KO groups. **g.** Food consumption ratio (Left) and water consumption ratios (Right) in the control and Eomes-KO groups. **h.** Left: the percentage of CD4 CTLs in the control (n=12) and Eomes-KO (n=14) groups. Right: flow cytometry analysis results showing the percentage of Granzyme B^+^ cells out of CD45^+^CD3^+^CD4^+^ cells in livers. **i.** Graphs showing the percentages of senescence-associated β-Galactosidase^+^ (SA-β-Gal) cells in non-immune liver cells (CD45^-^, Left) and immune cells (CD45^+^, Right) in control (n=12) and CD4 CTL-depleted (n=14) groups. **j**. Representative flow cytometry plots showing P16^+^P21^+^ cells out of CD45^-^ cells in the livers of control (Upper) and Eomes-KO (Lower) groups. **k.** Graphs showing the percentages of p16^Ink4a+^ and P21^+^ cells in non-immune liver cells (CD45^-^, Left) and immune cells (CD45^+^, Right) in control (n=6) and Eomes-KO (n=7) groups. Representative immunofluorescence images showing P16 and P21 immunolabeling in liver lobules from control (upper) and Eomes-KO mice (lower). Scale bar indicating 50 μm. **l.** Experimental setup: 16 months old control and Eomes-KO mice were subjected to a TMX regimen that included 3 days of IP TMX injection (100 µl) followed by an alternating dietary regimen of two weeks on TMX chow and two weeks on regular chow, lasting for a total of 40 weeks. **m.** Kaplan-Mayer survival curve for control (n=9) and Eomes-KO (n=9) mice. Censoring is indicated by the black live (|) mark. **n.** The change (%) in weight compared to baseline weight before the experiment (Y-axis) over 40 weeks (X-axis) in the control (n=6) and Eomes-KO (n=8) groups. Change in weight = (weight at week X/weight before the experiment) x 100. Bars indicate mean ± SEM from two (**a-i, l-n**) or one (**j-k**) independent experiments. Data were analyzed using two-tailed Student’s t-tests that were paired (**a**) or unpaired (**d, f-k, n**), or with log-rank (Mantel-Cox) tests (**m**). The exact P-values are presented in the figures. * = P value < 0.05.

To evaluate the effect of CD4 CTLs on senescent cell accumulation, livers and spleens were excised from the control and Eomes-KO mice and analyzed by flow cytometry (Methods). Analysis of the immune compartment revealed reduced frequencies of both CD4 CTLs and CD4 T cells expressing GzmB in the livers and spleens of Eomes-KO mice relative to control mice (Fig. 2h, Extended Data Fig.4a). Notably, no differences were observed in overall leukocyte infiltration of the liver (Extended Data Fig. 4b). Moreover, other CD4 T cell subsets in the liver and spleen did not differ between groups (Extended Data Fig. 4c-h). To assess senescence, we examined the liver tissue as an indicator for overall body senescence ^5,33^. First, we measured SA-β-Gal activity, a known marker for senescent cells ^11,34^. Both immune (CD45^+^ cells) and non-immune (CD45^-^ cells) cells within the livers had increased frequencies of SA-β-Gal^+^ cells in the Eomes-KO than in the control group (Fig. 2i). Furthermore, analysis of cells positive for two additional senescence markers (p16^ink4a+^p21^+^ cells) out of non-immune (CD45^-^) and immune (CD45^+^) cells, revealed increased frequencies of these cells in the Eomes-KO mice than in the control mice (Fig. 2j-k). Given the increased load of senescent cells in Eomes-KO mice we sought to test whether CD4 CTLs could affect lifespan. Eomes-KO and control mice (15 months old) were subjected to an alternating regimen of TMX-containing food and normal chow for 40 weeks, and their body weight and survival were assessed weekly (Fig. 2l, Methods). While the survival probability was estimated at 56% in the control group, it was estimated at 11% in the Eomes-KO group (Fig. 2m, p-value = 0.0262). In addition, 30 weeks after the initiation of the TMX regimen, a significant difference in body weight was observed between the groups (Fig. 2n), plausibly attributed to reduced activity and increased food consumption among these Eomes-KO mice. Collectively, these results indicate that CD4 CTLs limit senescent cell accumulation, a phenomenon that may support a causal link between CD4 CTL responses and longevity.

### CD4 CTLs locally differentiate in a mouse model of liver cirrhosis and mitigate cellular senescence and disease outcomes

In many respects, the microenvironment generated in the context of tissue aging is similar to that characteristic of long-lasting tissue inflammation in that SCs and fibrosis accumulate and promote tissue damage ^35,36^. Senescence has been implicated in the pathogenesis of various chronic immune-mediated diseases, including rheumatoid arthritis ^37^, pulmonary fibrosis ^38^, and liver cirrhosis ^39,40^. To evaluate the involvement of CD4 CTLs in a local model of chronic inflammation, we employed the carbon tetrachloride (CCL_4_)-induced liver cirrhosis model, which has been shown to entail fibrosis and SC accumulation ^39^. Young Eomes-KO (CD4-Cre^ERT2+/-^Eomes^fl/fl^) and control (CD4-Cre^ERT2-/-^Eomes^fl/fl^) mice (4-5 months old) were intraperitoneally (IP) injected with TMX for 3 days, followed by weekly alternating combined IP injections of TMX+CCL_4_ (CCL_4_-Control, CCL_4_-Eomes-KO) for 45 days (Methods). Another group of control mice was injected with the vehicle (TMX-Control). Mice were weighed weekly and then killed on day 45 for further analysis (Fig. 3a). Remarkably, flow cytometry analysis of liver and blood-derived CD4 T cells revealed increased frequencies of CD4 CTLs in the CCL_4_-Control compared with the TMX-Control group (Fig. 3b-c). As expected, no significant change in the frequency of CD4 CTLs was observed in the livers and blood of CCL_4_-Eomes-KO mice compared to those of TMX-Control animals (Fig.3b-c). Notably, CD4 CTLs did not accumulate in spleens of CCL_4_-Control mice, suggesting that their accumulation was primarily localized to the liver (Fig.3b). Analysis of other liver-derived CD4 T-cell subsets revealed markedly increased frequencies of Tregs (CD4^+^FOXP3^+^) in CCL_4_-Eomes-KO mice compared to both controls and increased frequencies of exhausted cells (CD4^+^CD44^+^PD1^+^) compared to TMX-Control mice (Fig.3d). Frequencies of liver effector memory CD4 T cells (CD4^+^CD44^+^CD62L^-^) were similarly reduced in both CCL_4_-Eomes-KO and CCL_4_-Control as compared to TMX-Control mice (Fig.3d). Furthermore, no differences in Treg, effector memory, or exhausted cells were observed between the groups in the blood or spleen (Extended Data Fig. 5a-b). Together, the signature of CD4 T-cell subsets in the livers of CCL_4_-Eomes-KO mice points to an immune environment that is reminiscent of fibrosis ^35^. Clinically, whereas weight changes were not observed throughout the experiment (Fig. 3e), livers excised from CCL_4_-Eomes-KO mice displayed more extensive scarring than those from CCL_4_-Control mice (Fig. 3f). Next, we measured the serum levels of aspartate transaminase (AST) and alanine transaminase (ALT), both indicators of liver damage ^41^. While both CCL_4_-Eomes-KO and CCL_4_-Control mice exhibited higher ALT levels, only the CCL_4_-Eomes-KO group showed elevated levels of AST compared to TMX-Control mice (Fig. 3g). Histological analysis revealed increased fibrosis in the CCL_4_-Eomes-KO group than in the CCL_4_-Control group, as indicated by Sirius Red staining (Fig. 3h-i). Additionally, in a double-blind setting, 35.7% of histological sections from the CCL_4_-Eomes-KO group scored for severe fibrosis (grade D), whereas only about 16.5% of the CCL_4_-Control group received the same score (Extended Data Fig. 5c). Analysis of liver senescence cells using SA-β-Gal^+^ indicated an increase in overall liver senescence in the CCL_4_-Eomes-KO group compared to both the TMX-control and CCL_4_-Control groups (Fig. 3j). These results were supported by a higher number of p16^ink4a+^p21^+^ cells, primarily observed around the liver sinusoids and liver capsule (Fig. 3k; Extended Data Fig. 6a-b), in the CCL_4_-Eomes-KO group. The quantification of the senescent load, defined as the ratio between the number of senescent cells and the circumference of lobular hepatic veins, showed an increase in p16^ink4a+^p21^+^ cells in the livers of CCL_4_-Eomes-KO than in the CCL_4_-Control mice (Fig. 3l). Collectively, these results underscore the pivotal role of CD4 CTLs in moderating tissue damage via limiting cellular senescence in chronic inflammatory conditions like liver cirrhosis.

**Figure 3.**
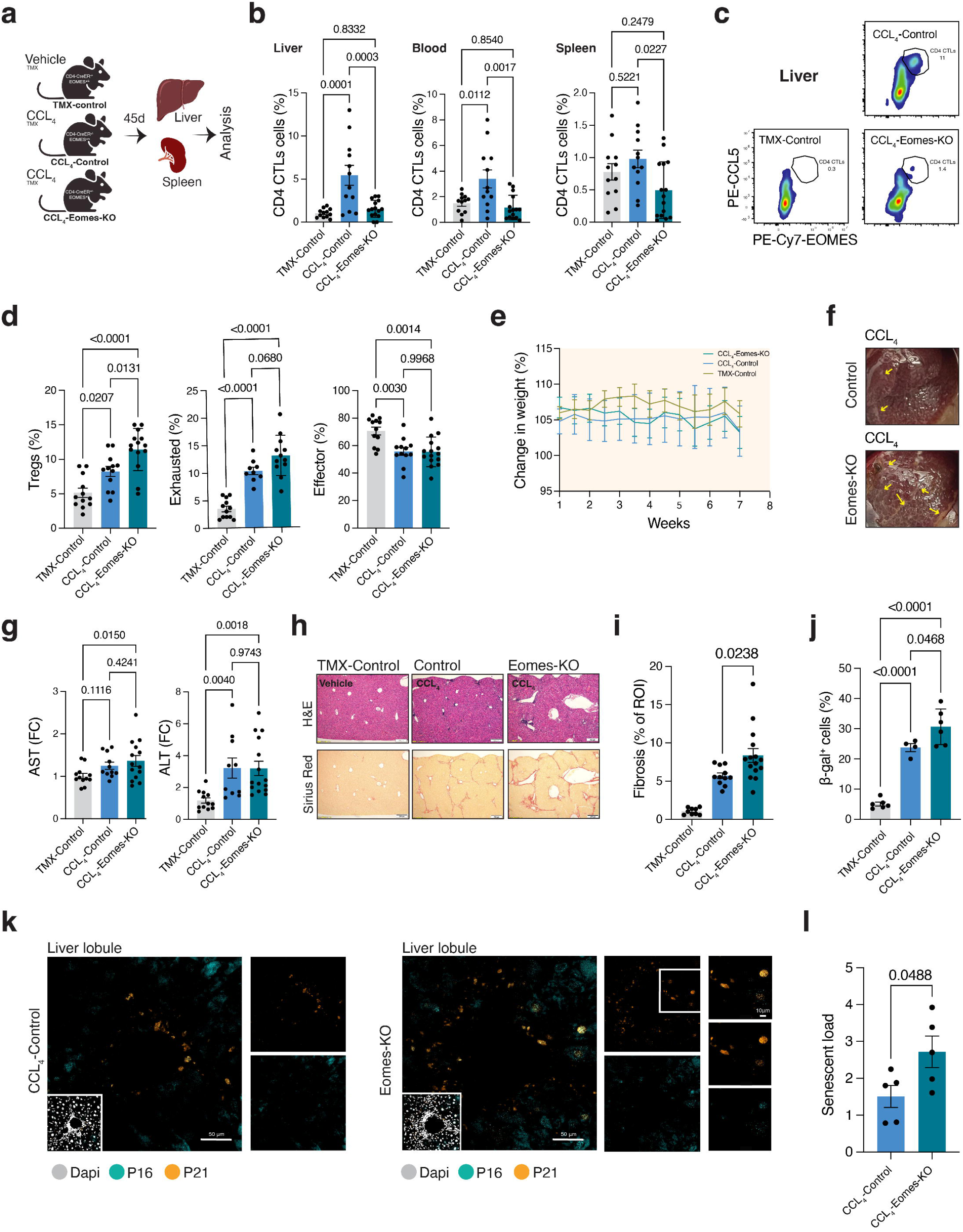
Depletion of CD4 CTLs is linked to increased fibrosis and senescence cell accumulation in a mouse model of CCL_4_-induced liver cirrhosis. **a.** Experimental setup: Cre^ERT2+/-^Eomes^fl/fl^ (CCL_4_-Eomes-KO) and Cre^ERT2-/-^Eomes^fl/fl^ (CCL_4_-Control) mice were subjected to a 45-day treatment with CCL_4_ + TMX to induce liver cirrhosis. Another group of Cre^ERT2-/-^ Eomes^fl/fl^ mice (TMX-Control) that received corn oil + TMX served as vehicle control. Spleens and liver were collected for further analysis after this treatment period. **b.** The percentages of CD4 CTLs in the livers (Left), blood (Middle), and spleen (Right) from TMX-Control (n=12), CCL_4_-Control (n=12) and CCL_4_-Eomes-KO (n=15) mice. **c.** Representative flow cytometry plots showing CD4 CTL percentages in the livers of TMX-Control, CCL_4_-Control, and CCL_4_-Eomes-KO mice. **d.** The percentages of Treg cells (CD45^+^CD3^+^CD4^+^FOXP3^+^, Left), exhausted cells (CD45^+^CD3^+^CD4^+^CD44^+^CD62L^-^PD1^+^, Middle), and effector cells (CD45^+^CD3^+^CD4^+^CD44^+^CD62L^-^PD1^-^, Right), and in the livers of TMX-Control (n=12), CCL_4_-Control (n=12), and CCL_4_-Eomes-KO mice (n=15) mice. **e.** The change (%) in weight compared to baseline weights before the experiment (Y-axis) over 7 weeks (X-axis) in the TMX-Control (n=11), CCL_4_-Control (n=12), and CCL_4_-Eomes-KO (n=12) groups. **f.** Representative images of CCL_4_-Control (Upper) and CCL_4_-Eomes-KO (Lower) mice demonstrating areas with liver scarring (arrows). **g.** Changes in AST (Left) and ALT (Right) measured by ELISA in the serum of TMX-Control (n=12), CCL_4_-Control (n=11), and CCL_4_-Eomes-KO (n=14) mice compared to the average level in the TMX-Control group. Each dot represents one mouse. **h.** Representative liver tissue images stained with H&E (Upper) and Sirus Red (Lower) for the TMX-Control, CCL_4_-Control, and CCL_4_-Eomes-KO groups. Scale bar represents 200 μm. **i.** Quantitative analysis of Sirus Red staining area calculated as the percentage of liver tissue area strained with Sirus Red out of all tissue area (ROI) for TMX-Control (n=9), CCL_4_-Control (n=11), and CCL_4_-Eomes-KO (n=15) mice. Each dot represents the average of at least two liver sections from one mouse. **j.** Graphs showing the percentages of senescence-associated β-Galactosidase^+^ (SA-β-Gal) cells out of total liver cells in TMX-control (n=6), CCL_4_-Control (n=4), and CCL_4_-Eomes-KO (n=6) groups. **k.** Representative immunofluorescence image showing P16 and P21 immunostaining in liver lobules. CCL_4_-Control (Left) and CCL_4_-Eomes-KO (Right) groups. Scale bars are indicated in the figures. **l.** Quantitative analysis of senescence load in the area of the lobular hepatic vein in the CCL_4_-Control group (n=5) and the CCL_4_-Eomes-KO group (n=5), calculated as the ratio between the number of senescent cells (P16^+^P21^+^) and hepatic vein circumference. Bars indicate mean ± SEM from one (**j-l**) or two (**a-i**) independent experiments. Data were analyzed using one-way ANOVAs with Tukey correction for multiple comparisons (**b, d, g, j**) or unpaired two-tailed Student’s t-tests (**i, l**). Exact P-values are presented in the figures.

## Discussion

Evidence increasingly links cellular senescence with the individual aging trajectory and age-related diseases ^42,43^. Whereas the immune system is central to clearing SCs and maintaining tissue homeostasis ^5,44^, the mechanisms through which the immune system modulates SCs remain elusive. Here, we found that a specific subset of CD4 T cells, known as CD4 CTLs, that accumulate with aging played a significant role in restricting cellular senescence and frailty. This observation was not limited to an aging environment, as when using an adult mouse model of liver cirrhosis, we observed that CD4 CTLs locally differentiated in the liver and impacted local senescent cell load and tissue pathology. Overall, we propose that CD4 CTLs differentiate in the context of tissue senescence, whereupon, in contrast to the suppressive functions of Treg and exhausted cells, they contribute to the clearance of SCs.

Our previous study revealed a gradual change in the CD4 T-cell landscape with age that mirrors the process of aging ^24^. CD4 CTLs accumulate relatively late in the process of aging and can occupy up to 50% of the CD4 T cells, as also observed in a Japanese cohort of supercentenarians ^27^. While these cells appear to mark a stage of aging, the signals promoting their differentiation and their role remain unclear. Here, by adaptively transferring CD4 T cells from young (CD45.1) to old (CD45.2) mice, we demonstrated that CD4 T cells can differentiate into CD4 CTLs in a SC-rich environment and that prior senolytic treatment halted such differentiation. This was further supported by our *in vitro* findings, where the co-culturing of CD4 T cells with senescent fibroblasts facilitated their differentiation to CTLs. Furthermore, after conditionally depleting CD4 CTLs in old mice (20 months old), we observed increased SC load along with a decline in physical performance metrics, including limb muscle endurance and overall physical activity. Long-term tracking of our model mice revealed a higher mortality rate in the absence of CD4 CTLs. Together, our findings suggest that CD4 CTLs can differentiate in a senescence-rich environment and undergo clonal expansion at the expense of other T cell subsets including naïve, exhausted, and Treg cells ^24^. At least in this context, they play a beneficial role by specifically restricting SCs ^30^. Their differentiation and function may, however, vary between individuals, possibly depending on genetic background such as the HLA haplotype, single nucleotide polymorphism, or other environmental factors such as chronic infection, and hence the individual’s capacity to mitigate cellular senescence.

Although CD4 CTLs accumulate in aging, they have also been observed within malignant tissues ^45^ or tissues subjected to infection or chronic inflammation ^45–47^. Except for the case of multiple sclerosis wherein these cells were associated with a more advanced stage of the disease ^48^, they primarily appear to play a beneficial role. Thus, to gain a general perspective on the role of CD4 CTLs as regulators of SCs, we utilized a mouse model of liver cirrhosis as a tissue-specific example of chronic inflammation. CD4 T cells in these mice, as expected, differentiated into CD4 CTLs during the 45 days of chronic liver inflammation. Furthermore, their conditional depletion resulted in increased frequencies of exhausted and regulatory T cells in the liver along with enhanced cellular senescence and worsened liver function and fibrosis. These results confirm that CD4 CTLs play a role in limiting the SC burden not only in the context of aging but also in tissues undergoing chronic inflammation and/or tumor growth. The unique contribution of CD4 CTLs within the realm of other cytotoxic effector functions (e.g., NK and CD8 T cells) should be further explored. At least one explanation may be related to their acting in the context of MHC class II-mediated antigen presentation, enabling them to target a broader set of antigens displayed by antigen-presenting cells and thus providing a more versatile mechanism for immune modulation and the mitigation of cellular senescence.

Collectively, these data illuminate a new biological role for CD4 T cells whereby they differentiate into CD4 CTLs as part of the immune effector functions involved in controlling cellular senescence. Our results showed that CD4 CTLs are central players involved in the maintenance of physiological homeostasis, offering potential avenues for therapeutic interventions aimed add addressing many age-associated diseases, treating conditions characterized by chronic inflammation, and prolonging health-span. Further research is required to unveil the signaling cues required for their differentiation, the main tissues where they accumulate with aging, and whether differences exist between individuals in the capacity of CD4 CTLs to differentiate and propagate.

## Methods

### Mice

Young (2-5 months) and old (20-24 months) groups were used in this study for several strains of mice including wild-type (WT) C57BL/6 (Strain#: 000664), CD4-CreER^T2^ (Strain#: 022356), Eomes-floxed (Strain#: 017293), and CD45.1 mice (strain#: 002014). CD4-CreER^T2+/-^ Eomes^fl/fl^ mice were used in the study, and littermates that were negative for Cre were used as controls. All strains were purchased from the Jackson Laboratory. Mice were housed and cared for under specific pathogen-free conditions in Ben-Gurion University’s animal facility. All animal-related experiments received approval from the Ben-Gurion University Animal Care and Use Committee.

### Tamoxifen regimen

For inducible gene knockout experiments, we employed different tamoxifen (TMX) treatment regimens or combinations based on the specific experimental design: a) Three consecutive days of intraperitoneal (IP) injections with 100 µl of TMX [Catalog: 13258, Cayman Chemical] diluted in corn oil (0.01 g/ml), followed by a three-day rest interval and another set of three daily injections; b) IP injections of 100 µl of TMX diluted in corn oil (0.01 g/ml) twice weekly for a duration of six weeks (12 total injections); or c) An alternating dietary regimen comprising two weeks on TMX chow (TEKLAD .130856TD) then by either one or two weeks on regular chow, repeating this cycle for up to 40 weeks.

### Tissue processing

Spleens were harvested and mashed into a 70-μm cell strainer. Lysis of red blood cells was performed using 300 μl of Ammonium-Chloride-Potassium (ACK) buffer for 1.5 minutes (Lonza, Basel Switzerland). Blood was collected in EDTA-coated tubes (MiniCollect, Greiner Bio-One) from euthanized mice via right atrial cardiac puncture. Red blood cells were then lysed using blood lysis buffer (BD Biosciences), and the remaining leukocytes were washed twice and counted. After right atrial puncture and blood collection, livers were harvested and processed using a Liver Dissociation Kit (Catalog# 130-105-807, Miltenyi Biotec) according to the manufacturer’s instructions.

### In vivo administration of Navitoclax (ABT-263)

The senolytic drug navitoclax (ABT-263) was given at a dose of 50 mg/kg body weight, or a vehicle (10% ethanol, 30% PEG 400, 60% Phosal50) was used. Administration was performed via oral gavage. The treatment regimen consisted of five consecutive days of treatment, followed by a one-week rest interval, and then another five consecutive days of gavage with this drug or vehicle control.

### Liver fibrosis model

Liver fibrosis was induced by the IP injection of a 1.0 ml/kg dose of CCL_4_ [Catalog: 289116-100ml Sigma] twice per week for six weeks (12 total injections of CCL_4_). CCL_4_ was diluted in corn oil at a 1:4 ratio.

### Survival analyses

CD4-Cre^ERT2+/-^Eomes^+/+^or CD4-Cre^ERT2-/-^Eomes^+/+^ male and female mice at the age of 16 months were IP injected 3 times with TMX diluted in corn oil (0.01 g/ml), then fed with TMX chow (TEKLAD .130856TD) and regular chow at rotating 2-week intervals (2 weeks on/2 weeks off) for 10 months. On a weekly basis, mice underwent clinical evaluation and their body weight was measured. Mice that developed severe health conditions such as tumors, physical restriction, or wounds were censored. Group differences in survival were analyzed using the log-rank (Mantel-Cox) test. Lifespan was measured in weeks for the final analysis. Body weight was calculated as follows:

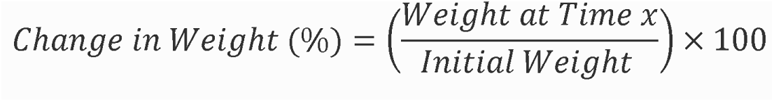

### Endurance Test

Mice used their forelimbs to suspend their body weight on a wire stretched between two posts 50□cm above the ground, and a pillow was used to prevent fall injuries. We evaluated the grip strength and endurance in each mouse with the wire-hanging test before and after TMX feeding. Mice were allowed to support themselves on a wire three times, the average duration they were able to maintain this position was normalized to body weight as mean endurance duration (sec)□×□body weight (g).

### Metabolic cages

For metabolic data acquisition, we used an 8-cage Promethion High-Definition Behavioral Phenotyping System (Sable Instruments, Inc., Las Vegas, NV, USA). Before experiments, mice were housed in the metabolic cage facility to allow them to adjust to the new environment for 7 days. Data were collected at 30-minute intervals for 84 hours of recording. Mice were maintained under a 12/12 h light/dark cycle (07:00–19:00) at an ambient temperature of ∼23°C. For data analysis, we assessed the following parameters: temperature, body mass, water and food consumption, movement distance and speed, and x and y breaks. Overall activity was defined as the sum of all distances traveled within the beam break system (x and y). This included fine movement (such as grooming and scratching) as well as direct locomotion. For analyses, we recorded each mouse before and after 45 days of TMX treatment. Then we calculated the performance ratio as follows: performance ratio = post-treatment parameter / pre-treatment parameter.

### Flow cytometry

For extracellular staining, cells were washed with FACS staining buffer (PBS containing 2% fetal bovine serum [FBS] and 1 mM EDTA) and incubated with an Fc receptor blocker (TrueStain fcX; BioLegend) for 5 minutes at 4℃. To differentiate between live and dead cells, the eFluor780-Fixable Viability Dye (eBioscience) was used according to the manufacturer’s instructions. Cells were incubated with primary antibodies for 25 minutes at 4℃ and then washed twice with FACS staining buffer. The following antibodies were used for surface staining: BUV395-conjugated anti-CD45.2 (104; BD Bioscience), BUV496-conjugated anti-CD4 (BD Bioscience), Brilliant Violet (BV) 421-conjugated anti-mouse CD3 (17A2; BioLegend), BV605-conjugated or BV510-conjugated anti-mouse CD279 (PD-1) (29F.1A12; BioLegend), AF700-conjugated anti-CD62L (Mel-14; eBioscience), BV650-conjugated anti-mouse CD44 (IM7; BioLegend), PerCP/cy5.5-conjugated anti-CD8 (53-6.7; BioLegend), and APC-conjugated anti-mouse CD45.1 (A20; BioLegend). After surface staining, intracellular labeling was performed. Cells were fixed and permeabilized using the Foxp3/Transcription Factor Staining Kit (eBioscience), blocked with rat serum (1 μl per 50 μl of staining buffer), and stained with the following antibodies: APC-conjugated anti-GzmB (QA16A02; BioLegend), PE-conjugated anti-CCL5 (2E9/CCL5; BioLegend), PE/cy7-conjugated anti-EOMES (Dan11mag; eBioscience), AF488-conjugated anti-FOXP3 (150D; BioLegend), AF647-conjugated anti-P21 (F-5, Santa Cruz Biotechnology) and CoraLite®488-conjugated anti-P16-INK4A (AG1328, Proteintech). For SA-β-gal staining, we used the CellEvent™ Senescence Green Flow Cytometry Assay Kit (eBioscience) according to the manufacturer’s instructions. All flow cytometry analyses were conducted using the CytoFLEX LX instrument (Beckman Coulter). Data analysis was performed with the FlowJo (v10.5.3) software. Gating strategies relied on fluorescence minus one controls, unstained samples, and unstimulated samples (when applicable). All samples in the experiment excluded dead cells, clumps, and debris.

### Immunohistochemistry

Livers were immersed in a 4% paraformaldehyde overnight at 4℃, then transferred into a 30% sucrose solution at 4℃ for 2 days and fixed in O.C.T Compound (Tissue-Tek). Sections (20-30 µm) of the liver were produced with a cryostat and kept at -20℃ in Glycerol. The sections were rinsed in a washing solution (0.05% Tween 20 in PBS), then permeabilized for 30 minutes in 0.5% Triton X-100 in PBS. Prior to staining, sections were incubated for 1 hour with a blocking solution containing 10% donkey serum in Antibody Diluting Buffer (Biomeda). Sections were then incubated for 48 hours with the following primary antibodies (1:100 in blocking solution): Rat anti-P21 (ab107099; Abcam), and Rabbit anti-CDKN2A/p16INK4a (EPR20418; Abcam). Sections were next rinsed 3 times in a washing solution and incubated for 1 hour with secondary antibodies conjugated to Alexa Fluor 546 (a11035; Invitrogen), Alexa Fluor 488 (a21208; Invitrogen), Alexa Fluor 488 (711545152; Jackson), Alexa Fluor 555 (ab150154; Abcam), and Alexa Fluor 633 (an-0082; Invitrogen) diluted 1:500 in PBS (or 1:250 for antibodies conjugated to Alexa Fluor 633). The sections were then rinsed in a washing solution twice before nuclei were stained with 4’,6-diamidino-2-phenylidole (DAPI; BioLegend). The sections were rinsed with washing solution one final time and then mounted on slides for examination under an Olympus FV1000 laser-scanning 4-channel confocal microscope (Olympus).

### Microscopy

Confocal images were produced with a 4-channel OLYMPUS XI81-ZDC confocal microscope. Images were acquired according to the sensitivity of the laser based on negative control sections of tissue or cell cultures. For image visualization and analysis, we used the Imaris software (Oxford Instruments) or ImageJ software (NIH).

### Senescent fibroblast co-culture

To establish primary cultures of lung fibroblasts, lung tissue from young (3-month-old) C57BL/6 mice was dissociated with the gentleMACS (Miltenyi Biotec) system and incubated at 37°C with Liberase™ for 1 hour, after which the tissue was washed and transferred into T175 flasks. Tissue samples were then incubated in Dulbecco’s Modified Eagle’s Medium (DMEM) (Gibco), supplemented with 10% v/v FBS (Invitrogen) and 1% Penicillin-Streptomycin at 37°C in a 5% CO_2_ incubator. Cells were allowed to migrate out from the tissue and grow to full confluence. Adherent cells were passaged by digestion with 0.025% Trypsin-EDTA when ∼90% confluent. After the second passage, cells were frozen at -80°C in FBS containing 10% DMSO, transferring 10^6^ cells into each Cryotube™ vial (Thermo Fisher). Experiments were performed with fibroblasts from individual cryovials of cells that were rapidly thawed in a 37°C water bath (1–2 min of agitation), resuspended in the culture media, and 10^5^ cells were then seeded into the separate wells of a 48-well plate for a 48-hour incubation at 37°C. Senescence was induced by treatment with Etoposide (Santa Cruz Biotechnology) dissolved in DMSO (30 mg/ml) and added to the culture medium (final concentration: 30 μM). Cells were treated with Etoposide for 48 hours, then washed with fresh medium and cultured for another six days. Control cells were thawed 72 hours prior to the initiation of the co-culture experiments. CD4 T cells were isolated from the spleens of young (2- to 3-month-old) mice spleen using a magnetic microbeads negative selection kit (EasySep Mouse CD4 T Cell Isolation Kit; STEMCELL Technologies) according to the manufacturer’s instructions. These CD4 T cells were activated with anti-CD3/anti-CD28 beads (Dynabeads, Gibco) in 96-well (U-shaped) plates (50 × 10^5^ cells per well) and transferred into the wells of the 48-well plate containing senescent or control fibroblasts. After 72 hours or 7 days, cells were collected and analyzed by flow cytometry. Cell supernatants were collected for ELISA analyses of cytokine levels. All cells were cultured at 37°C in a 5% CO_2_ incubator. Samples were analyzed in duplicate. All processing was performed using wide-bore pipette tips, and centrifugation speeds did not exceed 480 xg to minimize physical damage to the cells caused by shear forces.

### Histology

Tissues were fixed overnight in 4% paraformaldehyde and sent to Patho-Logica Ltd. (Israel) for subsequent tissue processing and clinical evaluation. Briefly, tissues were trimmed, placed in embedding cassettes, and processed for paraffin embedding. Sections (4 µm thick) were cut from the paraffin blocks, mounted on glass slides, and stained with hematoxylin and eosin (H&E) to assess pathological changes, and with Sirius Red (SR) to assess fibrosis. Imaging was conducted using an Olympus BX60 microscope (serial NO. 7D04032) with a DP73 camera (serial NO. OH05504).

### Fibrosis clinical scoring

Clinical scoring was conducted blindly by a pathologist (Patho-Logica Ltd, Israel). The degree of fibrosis measured by Sirius Red (SR) staining was evaluated under a light microscope at X4 magnification: Grade 0: No signs of fibrosis, Grade A: Very mild fibrosis, Grade B: Mild fibrosis, Grade C: Moderate fibrosis, Grade D: Severe fibrosis. For quantitative evaluation of the degree of fibrosis, we calculated the coverage area of SR staining out of a defined region of interest (ROI) consistent in size and shape for all tissue samples, analyzing at least two whole sections from each mouse using the ImageJ software.

### Liver Senescent Cell Load Calculation

To calculate the liver senescent cell load, we use ImageJ software to quantify the number of P16 and P21 positive cells around the hepatic vein within the liver lobules. For the calculation, we quantify at least 2 lobules hepatic veins from at least two sections for each mouse. The number of cells was normalized to the circumference of the lobule’s hepatic vein. The following equation was used for the analysis-

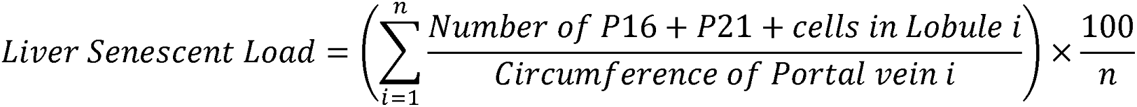

### ELISA

IL-2 and IFN-γ levels were measured with a sandwich ELISA kit (BioLegend) according to the manufacturer’s instructions. The levels of AST and ALT in murine serum were measured using EnzyChrom™ ELISA kits for AST and ALT (BioAssay Systems) according to the manufacturer’s instructions.

### Statistical analysis

Data are presented as the mean ± SEM unless otherwise stated. For statistical analyses, GraphPad Prism (v 10.0.3) was used. Paired t-tests were used for comparisons between multiple samples from the same biological samples. Log-rank (Mental-Cox) tests were used for survival analyses. For analyses of more than two groups, one-way ANOVAs were used with Bonferroni correction for multiple comparisons.

## Acknowledgments

We would like to thank Prof. A. Tarasiuk and Prof. A. Rudich for their guidance and support with the metabolic cages platform. We also appreciate Dr. M. Schiller for her comments on the manuscript.

## Authors contributions

Y.E. and A.M. conceived the project, designed experiments, and wrote the manuscript. I.F., N.P., A.Z., A.S., O.B., E.E., A.N., and K.R. performed experimental work and analyzed the data. L.R. and V.K. provided technical and scientific support.

