## Supplementary Information for "CD4 T Cells Acquire Cytotoxic Properties to Modulate Cellular Senescence and Aging"

**Supplementary Figure 1- Senescent cells induce $CD4^+$ CTL differentiation in vivo and in vitro.** **a.** Representative flow cytometry plots showing the gating strategy used to define CD4 T cell subsets. **b.** Quantitative analysis of CD4 T cells ($CD3^+CD4^+CD8^-$, Left) and naïve CD4 T cells ($CD3^+CD4^+CD44^-CD62L^+$, Right) out of transferred cells ($CD45.1^+$) in young or old WT mice (CD45.2). **c.** Representative images showing aged mice (22-24 months) in control (Left) or ABT-263 (Right) treated mice, demonstrating improved fur and health condition in the treated group. **d.** Representative images of P16 and P21 immunofluorescence staining for liver lobules of control (Upper) and ABT-263 (Lower) treated mice. Scale bar = 30µm (located in the upper right of the images). **e.** Left: A representative flow cytometry histogram plot showing the expression of EOMES in CD4 T cells harvested from the culture of control non-activated CD4 T cells (light blue), co-culture of activated CD4 T cells with control fibroblasts (dark blue), and co-culture of activated CD4 T cells with senescent fibroblasts (green), after 3 days of co-culture. Middle: Quantitative analysis of EOMES median fluorescence intensity (MFI). Right: percentage of granzyme $B^+$ cells out of activated CD4 T cells co-cultured with either control or senescent fibroblasts. **f.** Quantitative analysis of CD44 (Left) and PD1 (Right) MFIs. **(e-f)** Each dot represents an individual technical repeat. CD4 T cells were pulled and purified from 7 mice for each experiment. Bars indicate mean ± SEM from one **(b)** or two **(e-f)** independent experiments. Data were analyzed using a two-tailed Student's t-test **(b, e-f)** with exact P-values presented in the graphs.

**Supplementary Figure 2- Validation of CD4-Cre$^{ERT2+/-}$Eomes$^{fl/fl}$ mouse model for efficacy and specificity of EOMES deletion.** **a.** Spleen-derived T cells were activated using anti-CD3/anti-CD28 coated beads for 24 hours. Showing are flow cytometry analyses of $EOMES^+$ cells out of spleen-derived CD4 T cells purified from either CD4-CreERT2$^{+/-}$Eomes$^{fl/fl}$ mice (Eomes-KO; n = 5) or CD4-CreERT2$^{-/-}$Eomes$^{fl/fl}$ mice (Control; n = 5)) Left: percentage of $EOMES^+$ cells out of CD4 T cells. Middle: EOMES MFI in CD4 T cells of each group. Right panel: A representative flow cytometry histogram plot showing the expression of EOMES in non-activated (gray) or activated CD4 T cells purified from Control mice (green) or activated CD4 T cells purified from Eomes-KO mice (dark blue), after 24 hours in culture. **b.** Flow cytometry analysis of $EOMES^+$ cells out of spleen-derived CD8 T cells from Eomes-KO or Control after 24 hours of activation. Left: percentage of $EOMES^+$ cells out of CD8 T cells. Right: EOMES MFI in CD8 T cells. **c.** CD4 MFI in the $CD4^+$ T-cell population of Control and Eomes-KO mice. **d.** ELISA of IL-2 (Left) and IFNγ (Right) in the supernatants of CD4 T cells derived from Control (n=5) or Eomes-KO (n=5) mice and activated for 24 hours. Bars indicate mean ± SEM. Data were analyzed using a two-tailed Student's t-test, unpaired, with exact P-values presented in the graphs.

**Supplementary Figure 3- Control and Eomes-KO mice exhibit similar metabolic cage performance before tamoxifen administration. a.** Experimental Setup: 20-month-old Control (CreERT2$^{-/-}$Eomes$^{fl/fl}$) and Eomes-KO (CreERT2$^{+/-}$Eomes$^{fl/fl}$) mice were evaluated in metabolic cages for a week, after which TMX was administered IP. **b.** Representative temperature recording during the experiment. **c-g.** Representative comparison between Control (n=3) and Eomes-KO (n=3) groups for various parameters recorded over 312 hours at 30-minute intervals. These parameters include energy expenditure (kcal/hour; c), food consumption (grams; d), water consumption (grams; e), distance traveled (meters; f), and fine activity measured by x-axis beam breaks (g). **h.** The death rate in the Control group (30%) versus the Eomes-KO group (70%) during the time of both metabolic experiments (from the first to the second metabolic recording, a 45-day period).

**Supplementary Figure 4- Analysis of CD4 T-cell subsets in the liver and spleen of control and Eomes-KO mice. a.** The percentages of CD4 CTLs (Left) and CD4$^+$GzmB$^+$ cells (Right) in spleens of Control and Eomes-KO mice after the TMX regimen was administered. Regiment consisting of intraperitoneal injections of 100 µl of TMX for three days. This was followed by an alternating dietary regimen, where TMX chow and regular chow were alternated every two weeks, for a total duration of six weeks. **b-e.** The percentages of CD45$^+$ (**b**; Left), CD45$^+$CD3$^+$ (**b**; Right), CD3$^+$CD8$^+$ (**c**; Left), CD3$^+$CD4$^+$ (**c**; Right), naive (**d**; Left), effector (**d**; Right), Treg (**e**; Left), and exhausted (**e**; Right) cells in the livers of Control or Eomes-KO mice. **f-h.** Graphs showing the percentages of CD3$^+$ (**f**; Left), CD3$^+$CD4$^+$ (**f**; Right), Treg (**g**; Left), naive (**g**; Right), effector (**h**; Left), and exhausted (**h**; Right) cells in the spleens of Control or Eomes-KO mice. Subsets were defined as indicated in Supplementary Figure 1. Data were analyzed using a two-tailed, unpaired Student's t-test. The exact P-values are presented in the graphs.

**Supplementary Figure 5- Systemic immune response to CCL 4-induced liver inflammation in control and Eomes-KO mice. a-b.** The percentages of CD3$^+$CD4$^+$ (i), CD44$^+$CD62L$^-$PD1$^-$ effector cells (ii), CD3$^+$CD4$^+$FOXP3$^+$ Tregs (iii), and CD44$^+$CD62L$^-$PD1$^+$ exhausted cells (iv) in the blood (a) or in the spleen (b) of TMX-control (n=12), $CCL_4$ -control (n=12), $CCL_4$-Eomes-KO (n=15) mice. **c.** Pie plots showing the percentages of fibrosis severity scores (A-D) for $CCL_4$-Control (n=11), and $CCL_4$-Eomes-KO (n=15) mice Data were analyzed using one-way ANOVA, corrected by Tukey correction for multiple comparisons, with exact P-values presented in the graphs.

**Supplementary Figure 6- Characterization of senescent cells within the liver capsule and lobules in $CCL_4$-induced liver inflammation, comparing control and Eomes-KO mice a.** Representative immunofluorescence image showing P16 and P21 immunostaining of liver capsule for $CCl_4$-control (Upper) and $CCl_4$-Eomes-KO (Lower) groups. Scale bar indicating 50 μm. **b.** Representative immunofluorescence image showing P16 and P21 immunostaining in liver lobule in TMX-control group. Scale bar indicating 50 μm.

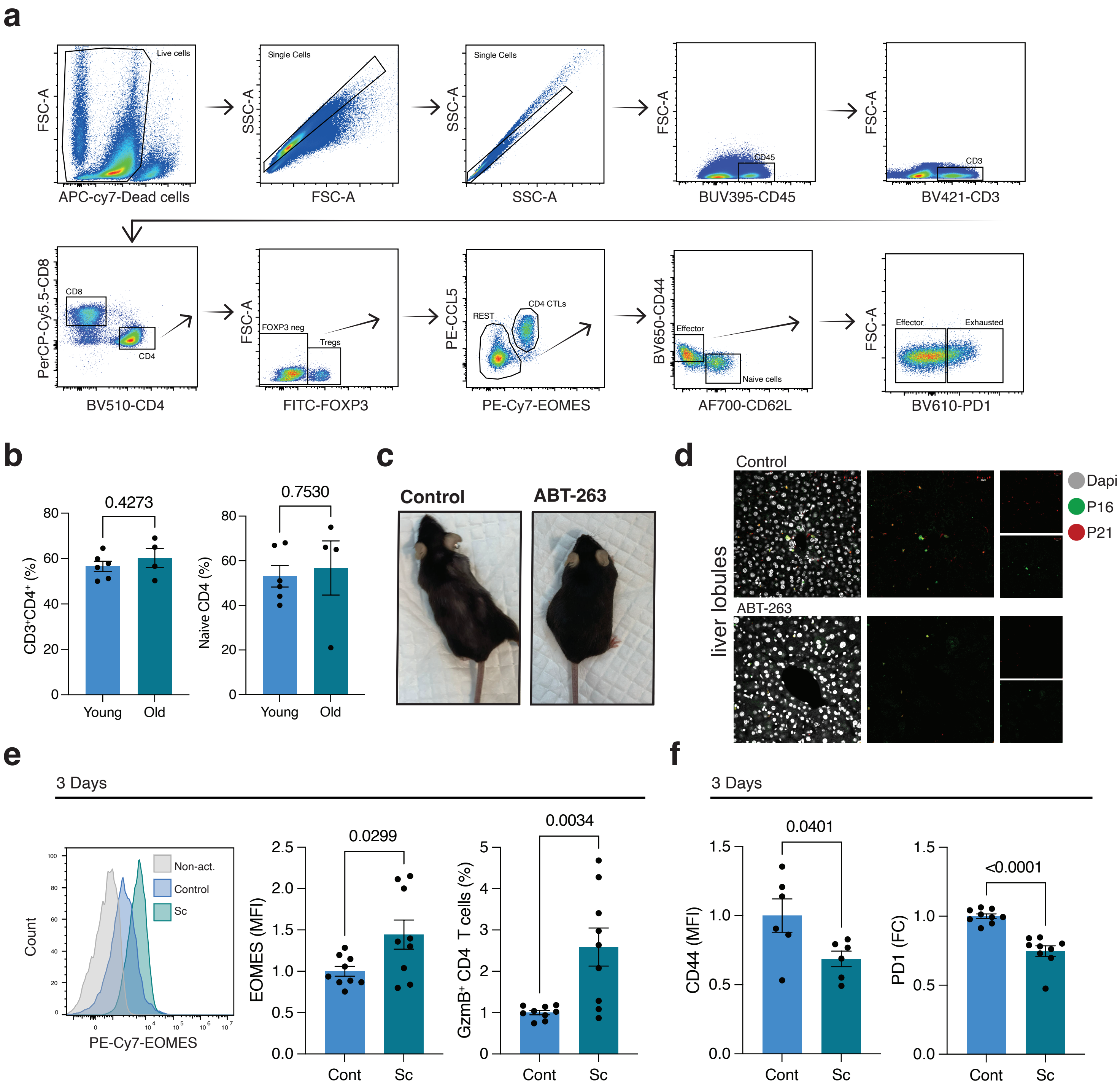


**Supplementary Figure 1**

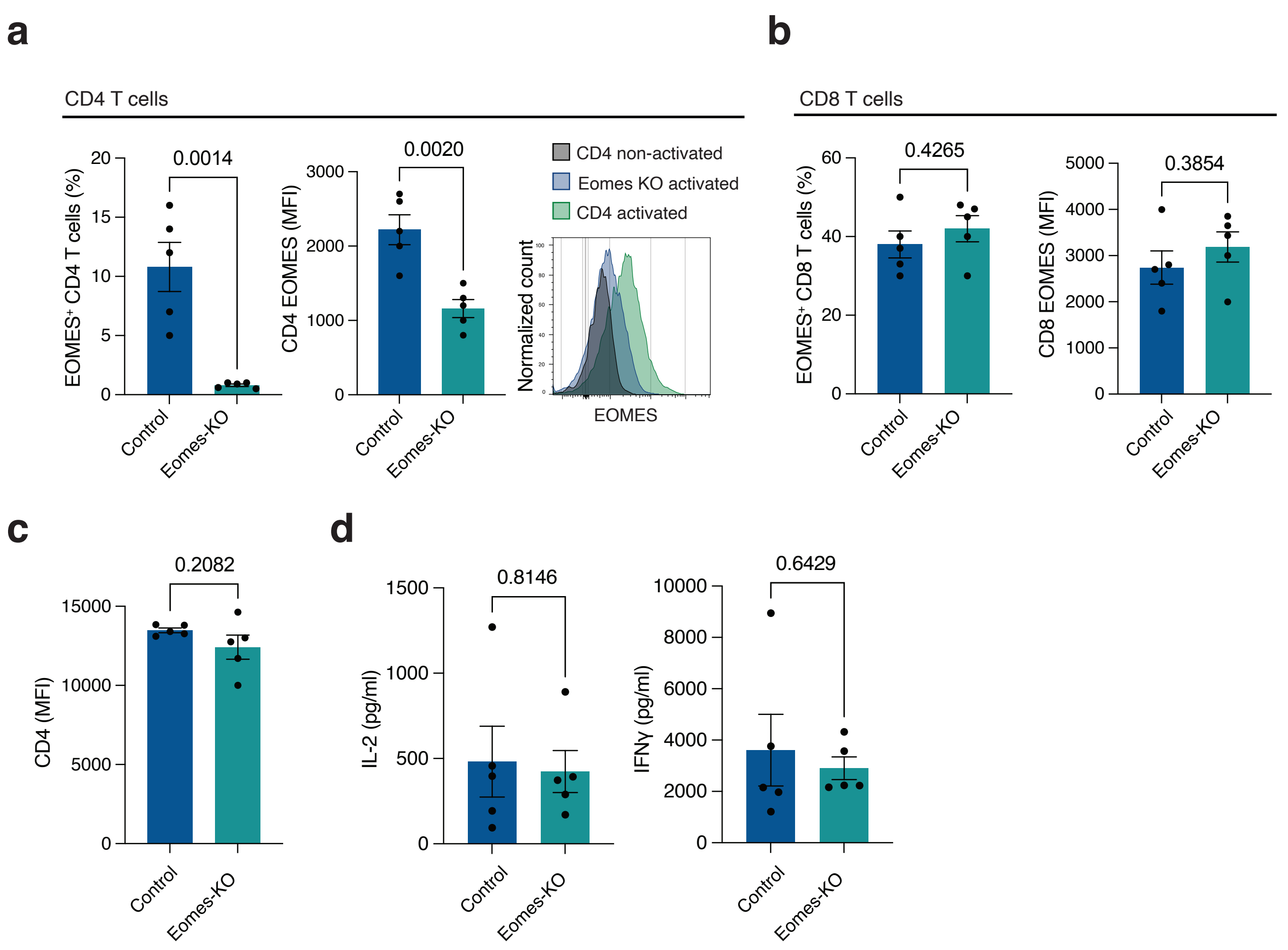


**Supplementary Figure 2**

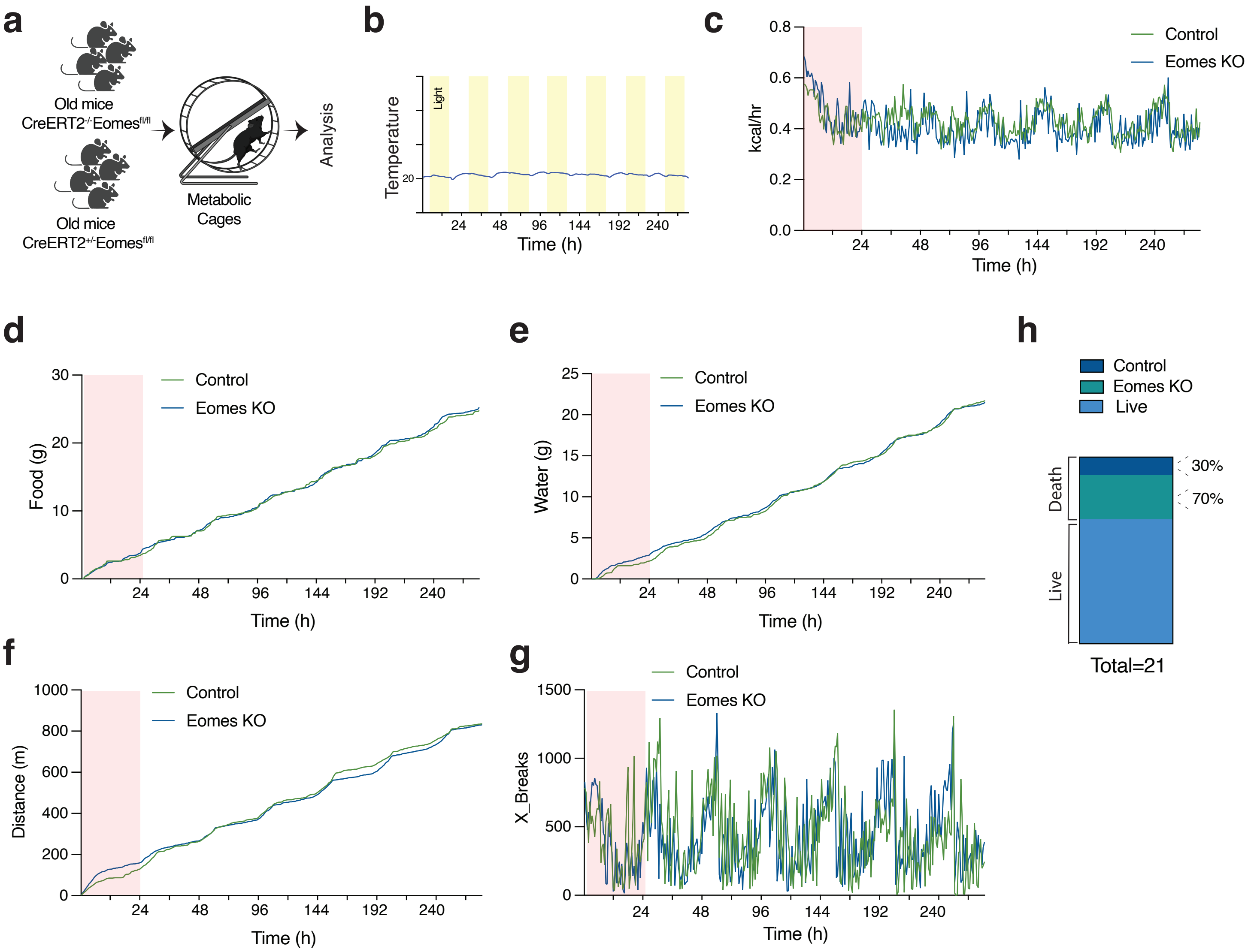


**Supplementary Figure 3**

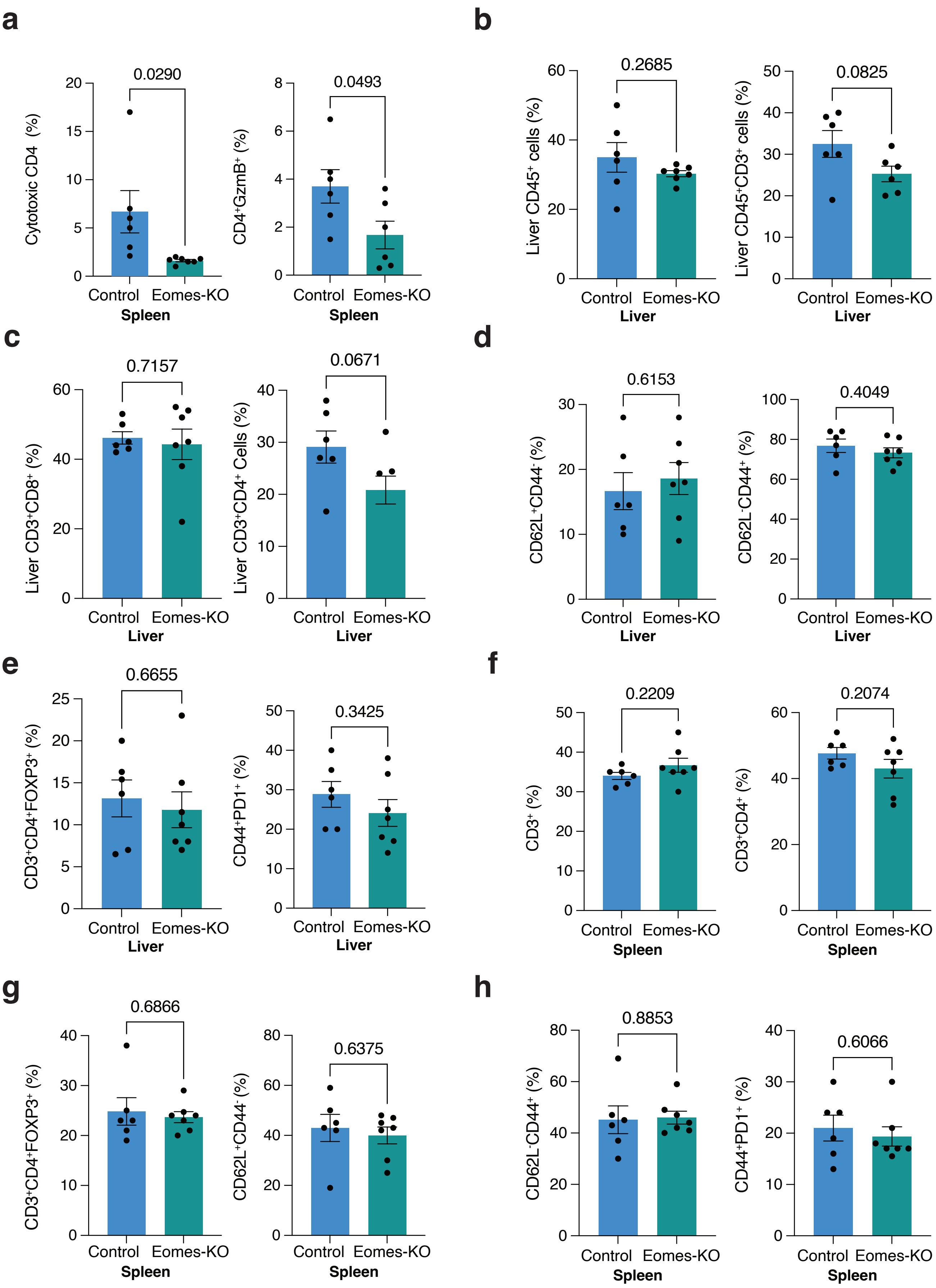


**Supplementary Figure 4**

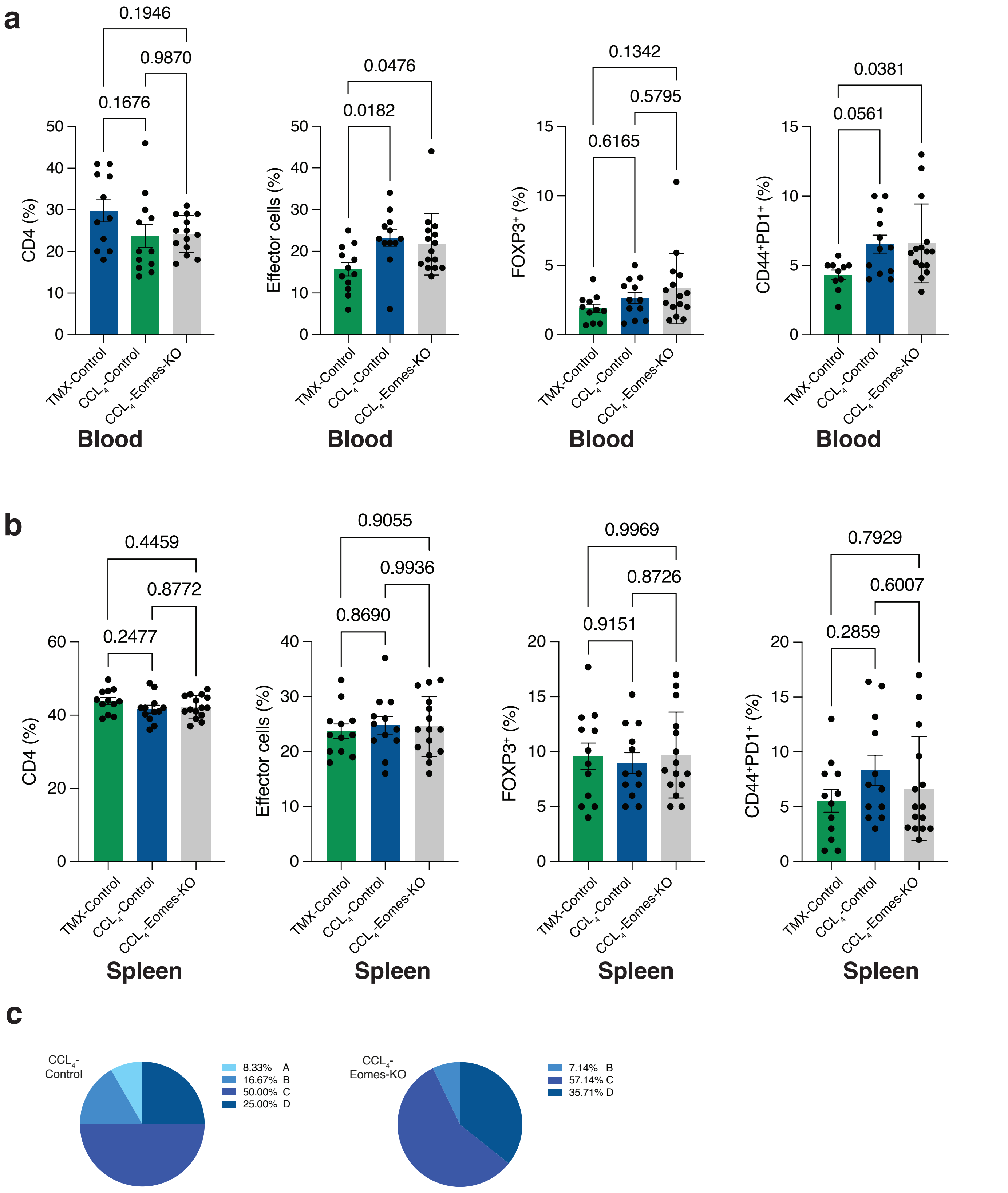


**Supplementary Figure 5**

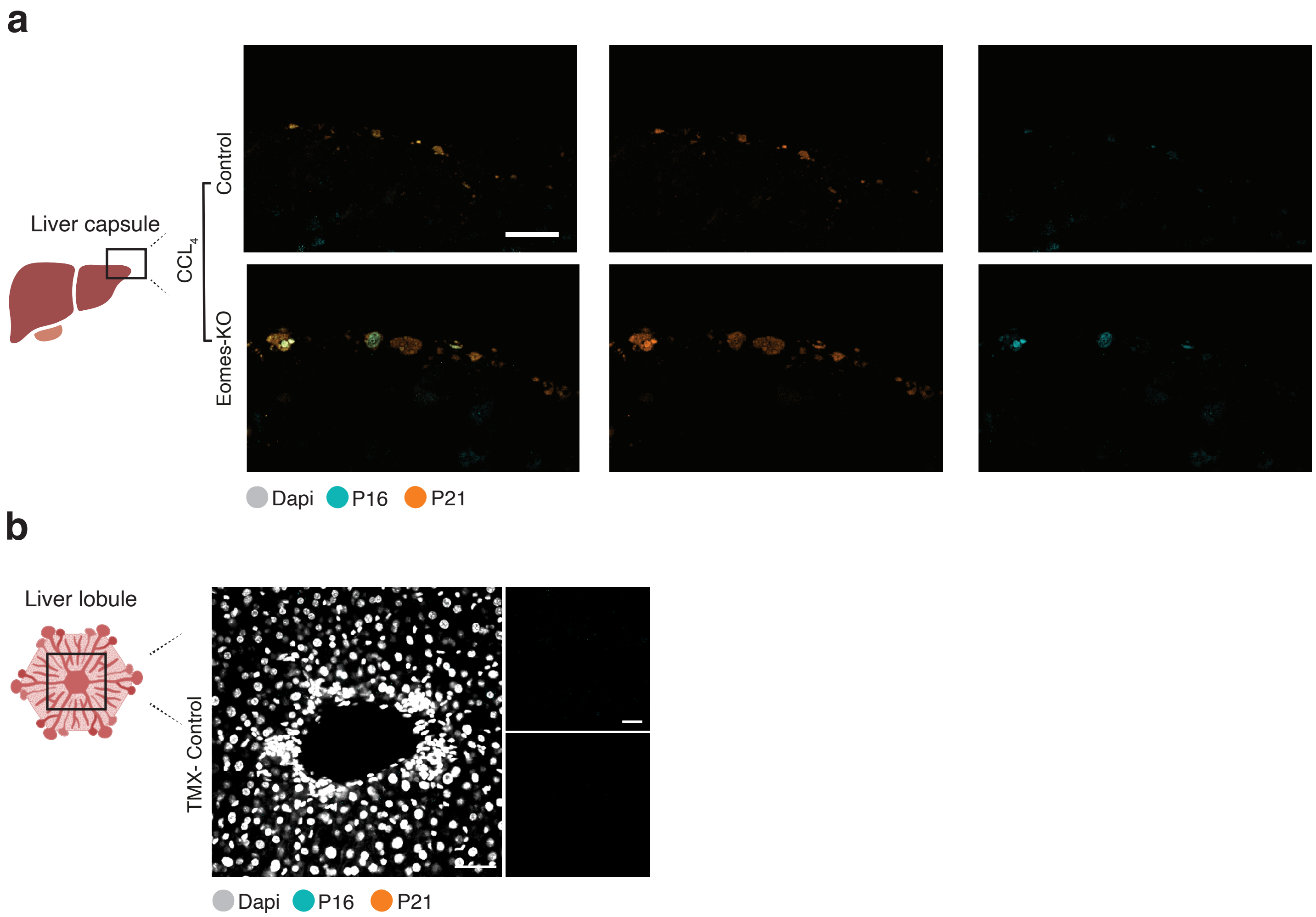


**Supplementary Figure 6**
